# Rapid whole genome sequence typing reveals multiple waves of SARS-CoV-2 spread

**DOI:** 10.1101/2020.06.08.139055

**Authors:** Ahmed M. Moustafa, Paul J. Planet

## Abstract

As the pandemic SARS-CoV-2 virus has spread globally its genome has diversified to an extent that distinct clones can now be recognized, tracked, and traced. Identifying clonal groups allows for assessment of geographic spread, transmission events, and identification of new or emerging strains that may be more virulent or more transmissible. Here we present a rapid, whole genome, allele-based method (GNUVID) for assigning sequence types to sequenced isolates of SARS-CoV-2 sequences. This sequence typing scheme can be updated with new genomic information extremely rapidly, making our technique continually adaptable as databases grow. We show that our method is consistent with phylogeny and recovers waves of expansion and replacement of sequence types/clonal complexes in different geographical locations.

GNUVID is available as a command line application (https://github.com/ahmedmagds/GNUVID).

## Introduction

Rapid sequencing of the SARS-CoV-2 pandemic virus has presented an unprecedented opportunity to track the evolution of the virus and to understand the emergence of a new pathogen in near-real time. During its explosive radiation and global spread, the virus has accumulated enough genomic diversity that we are now able to identify distinct lineages and track their spread in distinct geographic locations and over time [1-6]. Phylogenetic analyses in combination with rapidly growing databases [1, 7] have been instrumental in identifying distinct clades and tracing how they have spread across the globe, as well as estimating calendar dates for the emergence of certain clades [1-4]. This information is extremely useful in assessing the impact of early measures to combat spread as well as identifying missed opportunities [3]. Going forward whole genome sequences will be useful for identifying emerging clones or hotspots of reemergence.

In all of these efforts, identification of specific clones, clades, or lineages, is a critical first step, and there are few systems available to do this [1]. As of June 1^st^ there are already 35,291 and 4,636 complete genomes (>29,000bp) available at GISAID [7] and GenBank [8], respectively. To address the problem of identifying sequence types in SARS-CoV-2 and leverage these huge datasets, we took inspiration from a an approach used widely in bacterial nomenclature, multilocus sequence typing (MLST) [9]. Our panallelome approach to developing a whole genome (wgMLST) scheme for SARS-CoV-2 uses a modified version of our recently developed tool, WhatsGNU [10], to rapidly assign an allele number to each gene nucleotide sequence in the virus’s genome creating a sequence type (ST). The ST is codified as the sequence of allele numbers for each of the 10 genes in the viral genome.

Here we show that this approach allows us to link STs into clearly defined clonal complexes (CC) that are consistent with phylogeny. We show that assessment of STs and CCs agrees with multiple introductions of the virus in certain geographical locations. In addition, we use temporal assessment of STs/CCs to uncover waves of expansion and decline, and the apparent replacement of certain STs with emerging lineages in specific geographical locations.

## Results and Discussion

We developed the GNU-based Virus IDentification (GNUVID) system as a tool that automatically assigns a number to each unique allele of the 10 open reading frames (ORFs) of SARS-CoV-2 (**Figure 1A**) by modifying our tool WhatsGNU [10] (**Supplementary Methods**). GNUVID compressed the 104,220 ORFs in 10,422 high quality GISAID genomes (**Supplementary Table 1**) to 6244 unique alleles in less than one minute on a standard desktop, achieving 17-fold compression and losing no information. The majority of these alleles (65%) are for ORF1ab which represents 71% of the genome length (**Figure 1A**). Strikingly, the most abundant alleles of each ORF (except ORF1ab) were present in at least 79% of the 10,422 isolates, and for 8 ORFs (ORF3a-10) the allele that was observed in the earliest genomes was also the most prevalent, suggesting strong nucleotide level conservation over time.

**Figure 1.**
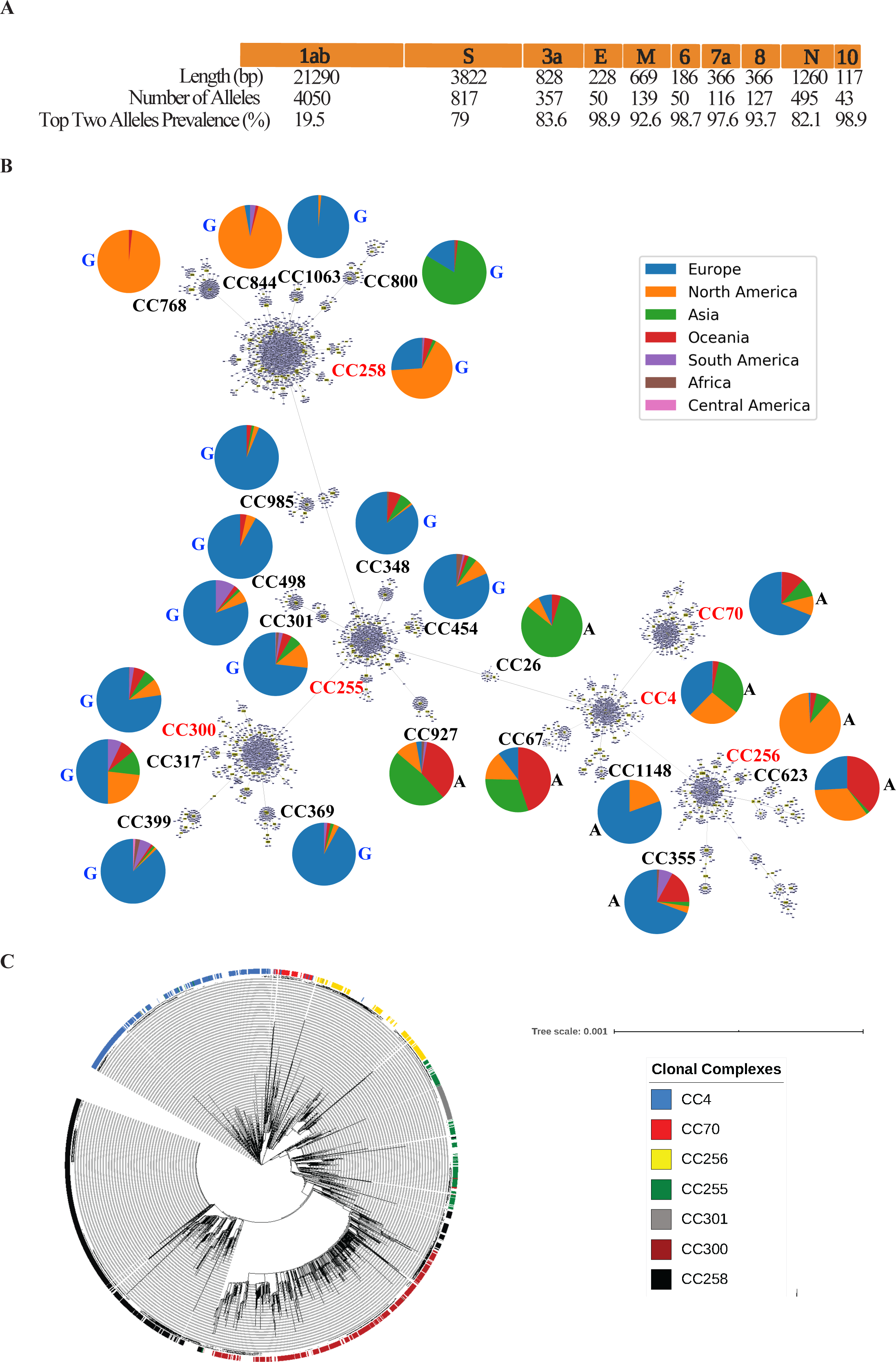
Sequence Typing Scheme for SARS-CoV-2. **A**. Map of SARS-CoV-2 virus genome showing the length in base pairs (bp) of the 10 ORFs, numbers of alleles in the current database, and the prevalence of the top two alleles of each ORF in the 10,422 database isolates. **B**. Minimum spanning tree from goeBURST of the 5510 Sequence Types (STs) showing the 24 Clonal Complexes (CCs) identified in the dataset. The largest six CCs are red and the other 18 CCs are in black. The pie charts show the percentage distribution of genomes from the different geographic regions in each CC. The letter A and G next to the pie charts represent the Spike ORF nucleotide at position 23403 in MN908947.3. The ancestral nucleotide is A and the mutation is G resulting in D614G amino acid change. At least 98% of the genomes of each CC had the reported nucleotide (except for CC26 where it was 93%). **C**. Maximum likelihood phylogeny of the 10,422 global high-quality SARS-CoV-2 sequences downloaded from the GISAID database (http://gisaid.org) on May 17^th^ 2020 (**Supplementary Table 1**). The tree is rooted on the reference sequence MN908947.3. The tree was visualized in iTOL. Only the most common seven CCs were shown for easier visualization. Nodes with 200-500 leaves were collapsed for better visualization. The raw tree is available as **Supplementary File 4**.

Some widespread alleles corresponded to mutations that have been hypothesized to be important to the evolution or pathogenesis of the virus. For instance, for the S gene, the gene for the Spike protein, 64% (526/817) of unique alleles have the A23403G (D614G) mutation (**Figure 1B**) that has been associated with the emergence of increased transmission whether through increased transmissibility [11] or lapses in control around this variant [3]. The first allele isolated and sequenced (allele 17) that carries this mutation was first recorded on January 24^th^ in China. The most common S gene allele that carries the A23403G mutation (allele 26) was present in 55% of the isolates. For ORF3a, which was shown to activate the NLRP3 inflammasome [12], 35% (126/357) of alleles have the G25563T (Q57H) mutation representing 33% of the isolates. The earliest sequenced, and most common, ORF3a allele that carries this mutation (allele 25) was isolated in France on February 21^st^ in a virus that also carries also the A23403G mutation in the spike gene.

To create an ST for each isolate GNUVID automatically assigned 5510 unique ST numbers based on their allelic profile (**Supplementary Table 2**). We then used a minimum spanning tree (MST) to group STs into larger taxonomic units, clonal complexes (CCs), which we define here as clusters of >20 STs that are single or double allele variants away from a “founder”. Using the goeBURST algorithm [13, 14] to build the MST and identify founders, we found 24 CCs representing 79% (4352/5510) of all unique STs (**Figure 1B**).

When the global region of origin for each genome sequence was mapped to each CC there was a strong association of some CCs with certain geographical locations. For instance, genomes from CCs 255, 300, 301, 317, 348, 355, 369, 399, 454, 498, 985, 1063, 1148 are predominantly from Europe while genomes from CCs 26, 800 and 927 are mainly from Asia (**Figure 1B**). Interestingly, genomes originating from the US appear to be associated with 2 very divergent CCs, potentially reflecting two major introductions. The first, CC256, is associated with locations on the West Coast, specifically Washington state. The first two isolates belonging to CC256 are from China followed by the first isolate from Washington (01/19/2020). The second predominant US CC, CC258, is closely related to other CCs found predominantly in Europe (**Figure 1B and 1C**). Isolates of CC258 were initially found and sequenced in Europe, followed by the US East Coast, and later in other US locations (**Figure 1B**). Interestingly, almost all isolates (99%) from CC258 and its descendants CCs 768, 800, 844 and 1063 (**Figure 1B**) carry the G25563T mutation in ORF3a, representing 88% of all isolates that carry this mutation; the other 12% are from STs that were not assigned CCs. CC800 is interesting for its geographic predominance in the Middle East (75% from Saudi Arabia and Turkey) and its close relationship to ST338 and ST258, which are mostly found in the US. This may signal a transmission event from the US to the Middle East.

To show that CCs are mostly consistent with whole genome phylogenetic trees we produced a maximum likelihood tree and mapped the CC designations onto the tree. Figure 1C shows that members of the same CC usually grouped together in clades (**Supplementary File 4**). One limitation of any ST/CC classification strategy is that paraphyletic groups can occur as a new ST arises from an older ST (e.g. CC301 emerged from CC255 making CC255 paraphyletic). While this means not all ST/CC groups will be monophyletic, this property of the nomenclature may be helpful in gauging emergence and replacement of an ancestral form.

To further validate our wgMLST classification system we compared it to the recently proposed “dynamic lineages nomenclature” for SARS-CoV-2 using the pangolin application[1]. A high percentage of viruses (90.5%;40-100%) with the same CC were assigned to the same lineage. When sublineages of the dominant lineage designation were included, this average rose to 99% (89-100%), showing strong agreement between these classification schemes (**Supplementary Table 2**).

Because we included collection dates for each genomic sequence, we can use STs and CCs to better understand the emergence and replacement of certain lineages in certain geographical regions over time. **Figure 2A** shows temporal plots of the most common 12 CCs around the world. This makes clear the emergence of new CCs over time such as CC255, CC300 and CC258. CC4, the earliest CC, started by representing 60% of sequenced genomes in mid-January, but had dropped to only 5% by mid-March.

**Figure 2.**
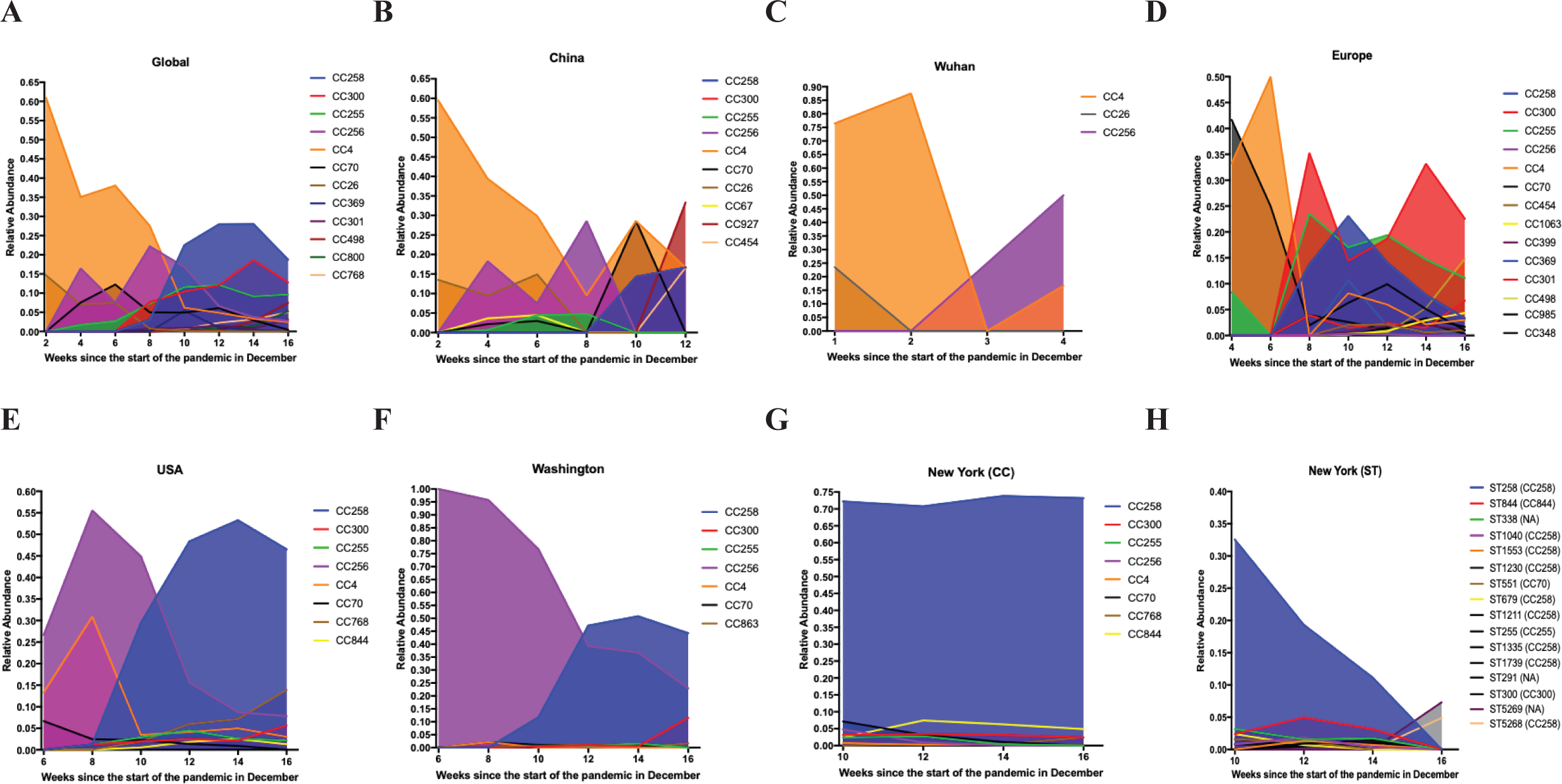
Temporal Plots of circulating STs/CCs at different geographical locations (Global, China, Wuhan, Europe, USA, Washington, NY (CC) and NY (ST)). The visualizations were limited to the most common CCs/STs. The raw data can be obtained from the authors upon request.

Of course, relative proportions of STs or CCs isolated and sequenced may be a highly biased statistic that is contingent upon where the isolate comes from, the decision to sequence its genome, and the local capacity to sequence a whole genome. Certain regions (US and Europe) clearly sequenced more genomes later in the pandemic compared to other countries.

Focusing on specific geographic regions may help to partially ameliorate this bias, and we chose to focus on three different regions (China, Europe and the US). The temporal plot of China shows expansion of local clones (CC4 and CC256) that likely spread to other countries early in the pandemic and then decreased in China over time. In contrast, two new CCs 927 and 454 appear to have emerged more recently with earliest isolation dates of March 18^th^ and April 16^th^, respectively, though this should be interpreted with caution because few sequences (n=7 and 6) were available/included. Interestingly, CC258 was first isolated in China in mid-March while it already represented 14% of the genomes in Europe by the end of February (**Figure 2B and D**), potentially reflecting transmission of new lineages back to China later in the pandemic. By the end of January, although CC4 represented 39% of the sequenced genomes in China, only one isolate (1/6) of CC4 was isolated in Wuhan, showing different patterns of circulating clones at the same timepoint in different parts of the same country (**Figure 2B and C**).

Interestingly, Europe showed a general CC diversity over time resembling that of the worldwide temporal plot, and then showed expansion of the local CC300 and CC255 after mid-February (**Figure 2D**).

The US plot (**Figure 2E**) reflects the two possible introductions on the west and east coasts from Asia and Europe, respectively, with the current dominance (more than 45%) of CC258. Focusing on Washington, it is interesting to note the possible replacement of CC256 by CC258 perhaps by introduction from the East Coast or Europe (**Figure 2F**) [2, 4]. In New York, a different pattern is seen with CC258 being persistently dominant (**Figure 2G**). However, a more granular view of STs in New York, not CCs, shows a shifting epidemiology with ST258 declining and the rise of closely related SLVs and DLVs of ST258 (**Figure 2H**).

While our wgMLST approach is rapid and robust it has several limitations. Because a change in any allele creates a new ST our method may accumulate and count “unnecessary” STs that have been seen only once or may be due to a sequencing error. This is partially ameliorated by the use of the CC definition that allows some variability amongst the members of a group. A large number of STs also may allow more granular approaches to tracking new lineages. Our method is also limited by the quality and extent of the database. For this implementation we limited the database to genomes that do not have any ambiguity or degenerate bases. However, these genomes could be queried through our tool to be assigned to the closest ST/CC. Another limitation is the stability of the classification system, some virus genomes may be reassigned to new CCs as clones expand epidemiologically, but this may also reflect a dynamic strength as circulating viruses emerge and replace older lineages.

## Conclusion

The genomic epidemiology of the 10,422 SARS-CoV-2 isolates studied here show six predominant CCs circulated/circulating globally. Our tool (GNUVID) allows for fast sequence typing and clustering of whole genome sequences in a rapidly changing pandemic. As illustrated above, this can be used to temporally track emerging clones or identify the likely origin of viruses. With stored metadata for each sequence on date of isolation, geography, and clinical presentation, new genomes could be matched almost instantaneously to their likely origins and potentially related clinical outcomes.

## Methods

All SARS-CoV-2 genomes (n=17,504) that are complete and high coverage were downloaded from GISAID [7] on May 17^th^ 2020. We kept 16,866 that were at least 29,000 bp in length and had less than 1% “N”s. Our wgMLST scheme was composed of all 10 ORFs in the SARS-CoV-2 genome [15]. The 10 ORFs were identified in the remaining 16,866 genomes using blastn [16] and any genome that had any ambiguity or degenerate bases (any base other than A,T,G and C) in the 10 open reading frames (ORF) was excluded. The remaining 10,422 genomes were fed to the GNUVID tool in a time order queue (first-collected to last-collected), which assigned a ST profile to each genome. The identified STs by GNUVID were fed into the PHYLOViZ tool [17] to identify CCs at the double locus variant (DLV) level using the goeBURST MST [13, 14]. CCs were mapped back to the STs using a custom script. Pie charts were plotted using a custom script. Temporal plots were extracted using a custom script and plotted in GraphPad Prism v7.0a.

To show the relationship between our typing scheme and phylogeny, we constructed a maximum likelihood tree. Briefly, we masked the 5’ and 3’ untranslated regions in the 10,422 genomes. We aligned these sequences using MAFFT’s FFT-NS-2 algorithm (options: --add --keeplength) [18] to the reference MN908947.3 [15]. A maximum likelihood tree using IQ-TREE 2 [19] was estimated using the HKY model of nucleotide substitution [20], default heuristic search options, and ultrafast bootstrapping with 1000 replicates [21]. The tree was rooted to MN908947.3. The tree and ST/CC data were visualized in iTOL [22]. We assigned a lineage [1] to each member of the 24 CCs using pangolin (https://github.com/hCoV-2019/pangolin) (options: -t 8). The GNUVID database will be updated weekly with new added high-quality genomes from GISAID [7]. Detailed methods are in **Additional file 1**.

## List of abbreviations

WhatsGNU: What is Gene Novelty Unit
GNUVID: Gene Novelty Unit-based Virus Identification
ST: Sequence Type
CC: Clonal Complex
SARS-CoV-2: Severe Acute Respiratory Syndrome Corona Virus 2
COVID-19: Corona Virus Disease 2019
MLST: Multilocus Sequence Typing
cgMLST: core genome MLST
wgMLST: whole genome MLST

## Additional files

**Additional file 1: Supplementary Methods** (txt, 34 Kb).

**Additional file 2: Table S1. Acknowledgment Table** (xls, 2.1 Mb).

**Additional file 3: Table S2. GNUVID Database Strains Report Table** (xlsx, 778 Kb).

**Additional file 4: Maximum Likelihood Tree of the 10422 strains** (nex, 369 Kb).

## Availability of data and material

The compressed genomes from our quality controlled dataset are available from the corresponding author and available online for download. The compressed database will be updated weekly on https://github.com/ahmedmagds/GNUVID. Source code for GNUVID can be found in its most up-to-date version here, https://github.com/ahmedmagds/GNUVID, under the GNU General Public License.

## Authors’ contributions

Conceptualization: AMM, PJP; Coding: AMM; Writing – Reviewing and Editing: AMM, PJP.

## Acknowledgements

We would like to thank Lidiya Denu and Michael Silverman for helpful comments and discussion. We would like to thank the Global Initiative on Sharing All Influenza Data (GISAID) and thousands of contributing laboratories for making the genomes publicly available. A full acknowledgements table is available as supplemental information.

## References

1. Rambaut A, Holmes EC, Hill V, O’Toole Á, McCrone JT, Ruis C, du Plessis L, Pybus OG: A dynamic nomenclature proposal for SARS-CoV-2 to assist genomic epidemiology. bioRxiv 2020:2020.2004.2017.046086.

2. Deng X, Gu W, Federman S, Du Plessis L, Pybus O, Faria N, Wang C, Yu G, Pan C-Y, Guevara H, et al: A Genomic Survey of SARS-CoV-2 Reveals Multiple Introductions into Northern California without a Predominant Lineage. medRxiv 2020:2020.2003.2027.20044925.

3. Worobey M, Pekar J, Larsen BB, Nelson MI, Hill V, Joy JB, Rambaut A, Suchard MA, Wertheim JO, Lemey P: The emergence of SARS-CoV-2 in Europe and the US. bioRxiv 2020:2020.2005.2021.109322.

4. Bedford T, Greninger AL, Roychoudhury P, Starita LM, Famulare M, Huang M-L, Nalla A, Pepper G, Reinhardt A, Xie H, et al: Cryptic transmission of SARS-CoV-2 in Washington State. medRxiv 2020:2020.2004.2002.20051417.

5. Shen L, Dien Bard J, Biegel JA, Judkins AR, Gai X: Comprehensive genome analysis of 6,000 USA SARS-CoV-2 isolates reveals haplotype signatures and localized transmission patterns by state and by country. medRxiv 2020:2020.2005.2023.20110452.

6. Chen Z-w, Li Z, Li H, Ren H, Hu P: Global genetic diversity patterns and transmissions of SARS-CoV-2. medRxiv 2020:2020.2005.2005.20091413.

7. Shu Y, McCauley J: GISAID: Global initiative on sharing all influenza data - from vision to reality. Euro Surveill 2017, 22.

8. Sayers EW, Cavanaugh M, Clark K, Ostell J, Pruitt KD, Karsch-Mizrachi I: GenBank. Nucleic Acids Res 2019, 47:D94–D99.

9. Maiden MC, Bygraves JA, Feil E, Morelli G, Russell JE, Urwin R, Zhang Q, Zhou J, Zurth K, Caugant DA, et al: Multilocus sequence typing: A portable approach to the identification of clones within populations of pathogenic microorganisms. PNAS 1998, 95:3140–3145.

10. Moustafa AM, Planet PJ: WhatsGNU: a tool for identifying proteomic novelty. Genome Biology 2020, 21:58.

11. Korber B, Fischer WM, Gnanakaran S, Yoon H, Theiler J, Abfalterer W, Foley B, Giorgi EE, Bhattacharya T, Parker MD, et al: Spike mutation pipeline reveals the emergence of a more transmissible form of SARS-CoV-2. bioRxiv 2020:2020.2004.2029.069054.

12. Siu K-L, Yuen K-S, Castaño-Rodriguez C, Ye Z-W, Yeung M-L, Fung S-Y, Yuan S, Chan C-P, Yuen K-Y, Enjuanes L, Jin D-Y: Severe acute respiratory syndrome coronavirus ORF3a protein activates the NLRP3 inflammasome by promoting TRAF3-dependent ubiquitination of ASC. The FASEB Journal 2019, 33:8865–8877.

13. Francisco AP, Bugalho M, Ramirez M, Carriço JA: Global optimal eBURST analysis of multilocus typing data using a graphic matroid approach. BMC Bioinformatics 2009, 10:152.

14. Feil EJ, Li BC, Aanensen DM, Hanage WP, Spratt BG: eBURST: Inferring Patterns of Evolutionary Descent among Clusters of Related Bacterial Genotypes from Multilocus Sequence Typing Data. Journal of Bacteriology 2004, 186:1518.

15. Wu F, Zhao S, Yu B, Chen YM, Wang W, Song ZG, Hu Y, Tao ZW, Tian JH, Pei YY, et al: A new coronavirus associated with human respiratory disease in China. Nature 2020, 579:265–269.

16. Altschul SF, Gish W, Miller W, Myers EW, Lipman DJ: Basic local alignment search tool. Journal of Molecular Biology 1990, 215:403–410.

17. Nascimento M, Sousa A, Ramirez M, Francisco AP, Carrico JA, Vaz C: PHYLOViZ 2.0: providing scalable data integration and visualization for multiple phylogenetic inference methods. Bioinformatics 2017, 33:128–129.

18. Katoh K, Misawa K, Kuma K, Miyata T: MAFFT: a novel method for rapid multiple sequence alignment based on fast Fourier transform. Nucleic Acids Res 2002, 30:3059–3066.

19. Minh BQ, Schmidt HA, Chernomor O, Schrempf D, Woodhams MD, von Haeseler A, Lanfear R: IQ-TREE 2: New Models and Efficient Methods for Phylogenetic Inference in the Genomic Era. Mol Biol Evol 2020, 37:1530–1534.

20. Hasegawa M, Kishino H, Yano T: Dating of the human-ape splitting by a molecular clock of mitochondrial DNA. J Mol Evol 1985, 22:160–174.

21. Hoang DT, Chernomor O, von Haeseler A, Minh BQ, Vinh LS: UFBoot2: Improving the Ultrafast Bootstrap Approximation. Mol Biol Evol 2018, 35:518–522.

22. Letunic I, Bork P: Interactive Tree Of Life (iTOL) v4: recent updates and new developments. Nucleic Acids Res 2019, 47:W256–W259.

