## Supplementary Methods for "Rapid whole genome sequence typing reveals multiple waves of SARS-CoV-2 spread"

### GNUVID: Supplementary Material

### 1 Installation

```
git clone https://github.com/ahmedmagds/GNUVID
cd GNUVID/bin
chmod +x *.py
pwd
export PATH=$PATH:/path/to/folder/having/GNUVID/bin
```

If it is needed permanently, the last line can be added to .bashrc or .bash_profile.

### 2 GNUVID.py Usage

#### 2.1 Dependency

- Python3 [1].
- Blastn [2].

#### 2.2 Input

1. database (precompressed (.txt) or a folder of individual genomes(.fna) to be compressed).
2. Reference File (MN908947.3_cds.fna).
3. Query CDS or whole genome FASTA file (.fna) or folder of query files.
4. Strains_order.txt: order of the strains by date of collection (**optional with -m but preferred**).
5. country_region.csv: Assigning regions (Europe, Asia..etc) to different countries (**optional with -m but preferred**).

#### 2.3 Command line options

*Create compressed database (ordered by date of collection and geographical regions assigned to the countries)*

```
GNUVID.py -m COVID19_10422_isolates/ -l Isolates_date_order.txt -cc country_region.csv -o GNUVID_results -p GNUVID MN908947.3_cds.fna CDS queries_folder/
```

Use precompressed database in Whole Genome Mode (the script will use blastn to identify the 10 ORFs in the WGS)

```
GNUVID.py -d db/GNUVID_05172020_comp_db.txt db/MN908947.3_cds.fna WG test_WG_query/
```

Use precompressed database in CDS Mode

```
GNUVID.py -d db/GNUVID_05172020_comp_db.txt db/MN908947.3_cds.fna CDS test_CDS_query/
```

#### 2.4 Output

*Always with -m or -d*

query_GNUVID_report.txt
Query_isolates_GNUVID_ST_Report.txt
GNUVID_date_time.log

*Always with -m*

prefix_comp_db.txt and prefix_DB_isolates_report.txt

### 3 Methods for the Precompressed Database

All SARS-CoV-2 genomes (n=17,504) that are complete and high coverage were downloaded from GISAID [3] on May 17th 2020. We kept 16,866 that were at least 29,000 bp in length and had less than 1% "N"s. The 10 ORFs were identified in the 16,866 genomes using blastn [4]. All isolates submitted before December 2019 were excluded.

```
GNUVID_FASTA_divider.py -l 29000 -N 1.0 COVID19_fasta gisaid_hcov-19_2020_05_17_05.fasta

for i in `cat /home/amoustafa/nas3/COVID-19/COVID19_11.list`;do blastn -task blastn -out COVID19_blast/${i}_results.txt -query MN908947.3_cds.fna -subject COVID19_fasta/$i.fna -evalue 0.000001 -outfmt '6 qseqid sseq sstart send pident qcovs'; done

Extract_fasta_sequence_blast_report.py COVID19_blast_extracted COVID19_blast/

GNUVID_database_customizer.py -p -i -l COVID19_cgMLST_10422_strains.csv COVID19_10422_strains COVID19_blast_extracted/

GNUVID.py -m COVID19_10422_strains/ -l COVID19_10422_strains _date_order.txt -o GNUVID_05172020 -p GNUVID_05172020 -cc country_continent.csv test_query/
```

The prefix_DB_isolates_report.txt was then used in PHYLOViZ tool [5] to identify CCs at the double locus variant (DLV) level using the goeBURST MST algorithm [6, 7]. CCs were mapped back to the STs using a custom script.

```
Clonal_complex_assigner.py -r resolve_10422.csv GNUVID_10422_strains_report_CC_assigned.txt phyloviz_eBURST_full_MST_10422.txt  prefix_DB_isolates_report.txt
```

### 4 References

1. **Python3: https://www.python.org/. Accessed 05 February 2019.**
2. Camacho C, Coulouris G, Avagyan V, Ma N, Papadopoulos J, Bealer K, Madden TL: **BLAST+: architecture and applications.** *BMC Bioinformatics* 2009, **10:**421.
3. Shu Y, McCauley J: **GISAID: Global initiative on sharing all influenza data - from vision to reality.** *Euro Surveill* 2017, **22**.
4. Altschul SF, Gish W, Miller W, Myers EW, Lipman DJ: **Basic local alignment search tool.** *Journal of Molecular Biology* 1990, **215:**403-410.
5. Nascimento M, Sousa A, Ramirez M, Francisco AP, Carrico JA, Vaz C: **PHYLOViZ 2.0: providing scalable data integration and visualization for multiple phylogenetic inference methods.** *Bioinformatics* 2017, **33:**128-129.

6. Francisco AP, Bugalho M, Ramirez M, Carriço JA: **Global optimal eBURST analysis of multilocus typing data using a graphic matroid approach.** *BMC Bioinformatics* 2009, **10:**152.
7. Feil EJ, Li BC, Aanensen DM, Hanage WP, Spratt BG: **eBURST: Inferring Patterns of Evolutionary Descent among Clusters of Related Bacterial Genotypes from Multilocus Sequence Typing Data.** *Journal of Bacteriology* 2004, **186:**1518.
